# Information structure of heterogeneous criticality in a fish school

**DOI:** 10.1101/2024.02.18.578833

**Authors:** Takayuki Niizato, Kotaro Sakamoto, Yoh-ichi Mototake, Hisashi Murakami, Takenori Tomaru

**Affiliations:** Department of Intelligent Interaction Technologies, Institute of Systems and Information Engineering, University of Tsukuba, Ibaraki; The Institute of Statistical Mathematics, Tokyo; Graduate School of Social Data Science, Hitotsubashi University, Tokyo; Faculty of Information and Human Science, Kyoto Institute of Technology, Kyoto

**Keywords:** Critical phenomena, Collective behaviour, Heterogeneous information process

## Abstract

The integrated information theory (IIT) measures the degree of consciousness in living organisms from an information-theoretic perspective. This theory can be extended to general systems such as those measuring criticality. Herein, we applied the IIT to actual collective behaviour (*Plecoglossus altivelis*). We found that the group integrity (i.e., Φ) could be maximised in the critical state and that several levels of criticalities existed in a group as subgroups. Furthermore, these fragmented critical groups coexisted with traditional criticality as a whole. The distribution of high-criticality subgroups was heterogeneous in terms of time and space. In particular, the core fish in the high-criticality group tended to be unaffected by internal and external stimulation, in contrast to those in the low-criticality group. The results of this study are consistent with previous interpretations of critical phenomena and provide a new interpretation of the detailed dynamics of an empirical critical state.

## Introduction

Critical phenomena are widely observed in living organisms.[1, 2, 3, 4]. The basic interpretation of a critical phenomenon is that the system is located midway between ordered and disordered states [5, 6, 1, 3, 7]. Although critical theoretical phenomena originate from statistical physics, they have several applications in living systems [8, 9, 10, 11]. Many researchers posit that living systems efficiently use these critical states, such as optimal information transfer[12, 13], high computational power[14, 15, 9], and inducing adaptive behaviour [16, 17, 18]. Furthermore, certain researchers have proposed building artificial critical systems using its high computational power, such as robots and computers [19, 11].

The criticality observed in nature, which we call empirical criticality in this paper, has been interpreted and developed based on theoretical criticality. However, applying theoretical criticality to empirical criticality has certain limitations. The main problem is that we cannot determine whether the fluctuations in the system arise from an external or internal cause because the actual system is finite and open to the environment [20]. This problem is important because the criticality of the system is derived from fluctuations therein. In the theoretical context, many researchers assume that the fluctuations in the system are relatively independent [8, 9, 21]. In other words, individual fluctuation is determined without any explicit rules [5]. Assuming independence, we can treat the fluctuation as the parameter determining criticality [9, 5, 22, 23]. This parameter tuning aspect enables many researchers to have insisted the criticality as the border between order and disorder. However, at the same time, the parameter tuning also implies that the system element cannot regulate its fluctuations. Consequently, the origin of criticality in nature becomes an essential problem because nature tunes this optimal parameter for the system. Although Bak et al. proposed the concept of ‘self-organised criticality’, no consensus exists among researchers regarding the optimal tuning[1, 24, 6, 21].

In contrast, many observations has revealed that the fluctuation is not simple random phenomena. The fluctuation that originates within an individual’s decision/stimulus selection in a system can be inherently controlled by the agent[25, 26, 27, 28]. This controllability enables the system to provide appropriate options for unpredictable events[29, 30, 31]. Thus, individual fluctuations are not uniform in time or space[30, 32] since this fluctuation depends on the individual’s situation. In addition to this problem, the uncertain boundary conditions of the system result in a complex behaviour [33]. These constraints on the inherent fluctuations hinder a standard analysis of the pure theoretical criticality of the system. As an incomplete observer, we cannot discriminate between the structural differences in inherent and external fluctuations [7].

The integrated information theory (IIT) was proposed to define human consciousness mathematically [34, 35, 36, 37, 7]. The basic concept of IIT is summarised in the proposition that the degree of consciousness can be measured as the difference between the entire system and the sum of its parts (i.e., Φ). Although IIT has its roots in neurological interest, its applications can extend to general systems [39, 40, 41, 42, 43, 44, 7]. For instance, as a purely theoretical application, the degree of integrity (Φ) accurately measures a critical state [46], such as the edge of chaos[41]. Niizato et al. also applied IIT to an empirical system (e.g., a fish school[43, 44] and body information[45]), and they found a characteristic structure that was not observed in other information-theoretic analyses.

The advantages of applying IIT to a living system are twofold. First, as discussed, the system integrity (i.e., Φ) measures the degree of criticality. We can examine the information structure of the system criticality using IIT. Second, the IIT does not discriminate between internal and external noise [36, 38, 47]. This attitude (indistinguishess between internal and external noise) is based on a paradigm from ‘what the system does’ (i.e., the system as an input–output function) to ‘what the system is’ (i.e., the system as the ability to do something) [47]. Therefore, the problem of indistinguishability between internal and external fluctuations gives suitable motivation to applying IIT analysis.

The second property has several implications for analysing living systems as following. If we ignore external and internal fluctuations, a stimulus from the outside cannot be discriminated from a stimulus from the inside. This property results in ‘finding the outside in the system’; that is, the entire system can be decomposed into certain subsystems (i.e., main complexes. See Methods), which are irreducible. These irreducible subsystems facilitate the analysis of the system on a mesoscopic scale (not individuals [48, 49], not the entire system [12, 25]). On this mesoscopic scale, we can examine more detailed structures within the empirical criticality.

This study aimed to reveal the essential factor for empirical criticality, which has not been confirmed in theoretical criticality, considering animal collective behaviour. We applied the IIT to real fish data (*Plecoglossus altivelis)*. We used two variables for IIT computation: orientation and speed, which are critical properties[12]. Throughout this study, we assumed that the group integrity (i.e. Φ) represented the degree of the critical state. The validity of these identifications is discussed in the model comparison Section (e.g., compared with the self-propelled particle model [50, 5] and Boids [51]). By applying the IIT to collective animal behaviour, we can comprehend the information process of criticality (i.e., the relationship between collective states and their information structure). The IIT analysis confirmed the traditional argument on criticality and a new aspect of criticality, defined as local criticality, which can coexist with traditional criticality.

## Result

### Brief review of IIT

First, we explain certain IIT concepts for readers unfamiliar with the IIT (we used IIT version 2.0, as four versions have been proposed). Although many concepts exist in the IIT, we focused on three aspects: the minimum information partition (MIP), main complexes, and integrated information (Φ).

As discussed in the Introduction, the major issue of the IIT concerns intrinsic information; it depends only on inner variables. No external system exists. This inner variable also changes in response to external and internal stimuli. Typical information theories in biology assume a relationship between external inputs and results. In this setting, observer interest focuses on the input–output relationship via the system of interest (i.e., ‘what the system does’). By contrast, IIT focuses on the system dynamics (i.e., ‘what the system is’): the mutual information between the past and present states. Let *X*^*t*^ be an internal variable vector (*X* = *X*_1_, *X*_2_, …., *X*_*n*_ ) at time *t*. The mutual information is then expressed as follows:

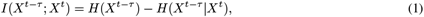

where *H*(*X*^*t*−*τ*^ ) is the entropy of the past states and *H*(*X*^*t*−*τ*^ |*X*^*t*^) is the conditional entropy of the past states given the present states.

However, mutual information contains information irrelevant for integrity, such as event correlation and historical effects. The IIT applies the MIP to the system to eliminate such an effect from this information. We must search for the weakest link among all possible cuts, which grows exponentially as the system size increases (this computational complexity forces us to limit the application of the IIT to a small system). In this study, we used the mismatch decoding method proposed by Oizumi et al., which has a relatively high computational speed. System integrity in the Φ^*^ is expressed as

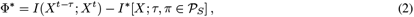

where *I*(*X*^*t*−*τ*^ ; *X*^*t*^)) is the mutual information between the current *X*^*t*^) and past states *X*^*t*−*τ*^, *S* is the set of all nodes of a given system, 𝒫_*S*_ is the set of all bi-partitions (total 2^|*S*|^ −1 partitions), *π* is an element of a set 𝒫_*S*_, and *I*^*^(*X*; *τ, π* ∈ *𝒫*_*S*_) is a ‘hypothetical’ mutual information, indicating the mismatched decoding in the partitioned probability distribution by *π*. More precisely, *I*^*^(*X*; *τ, π* ∈𝒫 _*S*_) is expressed as the partition max_*β*_ *Ĩ*(*β*; *X, τ*, _*S*_) that minimises Φ^*^ (see listed studies[38, 40] for further details about this expression).

Because Φ^*^ depends on partition *π*(∈ P_*S*_), the MIP is the partition that minimises the integrated information.

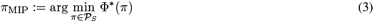

The integrated information for *π*_MIP_ is expressed as Φ^*^(*π*_MIP_). We denote Φ^*^(*π*_MIP_) Φ_MIP_. Notably, if Φ_MIP_ is equal to zero, the parts of the system are mutually independent; that is, there is no interaction between the parts. In this sense, Φ_MIP_ characterises the irreducibility of a system into its parts.

### Main complexes

In general, we can compute Φ_MIP_ for any subsystem in the system and not only for set *S*. We denote each Φ_MIP_ for subsystem *T* as 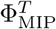, where *T* ⊂ *S*. A ‘complex’ is a subsystem *C*(⊂ *S*), where 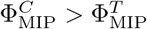 for all supersets of *S*. Note that the entire set *S* always satisfies complex conditions.

Based on this definition, we define the main complexes as those with a local maximum 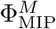.

### Definition (Main complex)

A main complex is a complex *M* satisfying 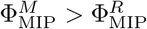 for the subsystem *R* ⊂ *M* .

This definition states that if two main complexes exist (say, *A* and *B*), then they are exclusive. IIT researchers consider these complexes to be the information core of the system, which may be related to our conscious experience. Although such information cores play a vital role in living systems, the validity of this assumption remains to be demonstrated [39, 43].

The main complex is among the most critical concepts of IIT. The main complex is a subset with a maximum Φ_MIP_. Based on its definition, several main complexes exist in a system. Tononi et al. suggested that the main complex, having a maximum Φ_MIP_, can serve as a consciousness based on their exclusive principle. In the present study, we do not insist that the Φ_MIP_ values correspond to the consciousness of the fish group; instead, we apply them as a measure of the system integrity, particularly the degree of criticality.

### Application of the IIT to a fish school

#### Data

We tracked the trajectories of *ayu* fish schools with *N* = 10 using seven samples (8–12 min recording length). Because the frame rate was 1/20 s, there were approximately 12,000 frames for the position data (i.e. ***x***(*t*) = (*x*_*i*_(*t*), *y*_*i*_(*t*))).

#### Variable settings for IIT 2.0

We applied IIT 2.0 to the direction of the fish (i.e., turning rate) and speed (i.e., acceleration). We computed each velocity vector ***v***_*i*_(*t*) from the obtained positional information, ***x***_*i*_(*t*), that is, ***v***_*i*_(*t*) = ***x***_*i*_(*t*) −***x***_*i*_(*t* −Δ*t*), where Δ*t* was 1/20 s. The turning rate, *dθ*_*i*_(*t*) is 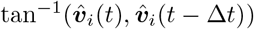, where hut is the unit vector of ***v***. The acceleration *ds*_*i*_(*t*) is ||***v***_*i*_(*t*)) −***v***_*i*_(*t* −Δ*t*)|| . Therefore, the group vector information is expressed as ***dθ***(*t*) = [*dθ*_1_(*t*), *dθ*_2_(*t*), …, *dθ*_10_(*t*)], ***ds***(*t*) = [*ds*_1_(*t*), *ds*_2_(*t*), …, *ds*_10_(*t*)]. Each Φ_MIP_ was computed from the time series of these vectors (Figure 1A). We set the maximum interval (*T*_max_) for the analysis. The vector [***dθ***(*t*), ***dθ***(*t* − Δ*t*), …, ***dθ***(*t* − *T*_max_Δ*t*)] was set for the analysis. We calculated Φ_MIP_ for three conditions: *T*_max_ = {200, 400, 600}, that is, 10, 20, and 30 s.

**Figure 1:**
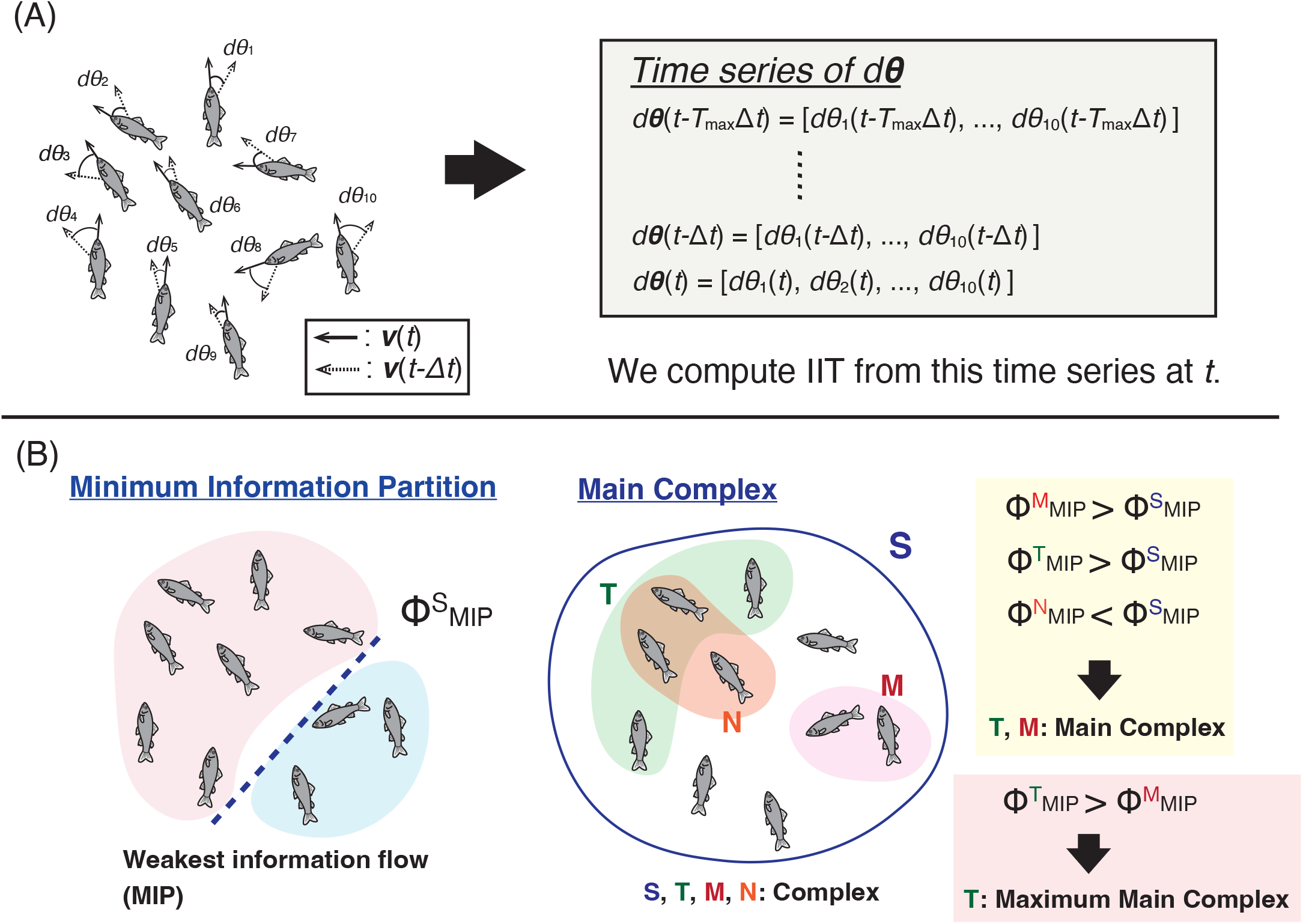
IIT computation methods and concepts. (A) Constructing *d****θ*** for IIT computation. We used the same method for *d****v***. (B) IIT concepts. (1) Minimum information partition (MIP): MIP-cut divides the fish school into two halves (coloured red and blue), which show the weakest information flow. The loss of information induced this cut implies Φ_MIP_. In this study, we set Δ*t* = 0.15 and *T*_max_ = 400. (2) Main complex: the maximum subset *T* has the maximum 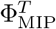 among any complexes (superset). In this case, *T* (green) and *M* (red) are the main complexes. In particular, the maximum main complex is given as *T* (green).

#### Setting a parameter for IIT 2.0 application

Oizumi et al. proposed approximation methods for the computational problem of the IIT [38, 52, 53]. In this study, we applied their ‘Practical Φ Toolbox for MATLAB’ to the fish data. An exhaustive method (computing all the possible MIPs) could be applied within a realistic computation time for such a small system.

Because the time delay *τ* (IIT has only two constraints: partition *π* and time delay *τ* ) was chosen as a suitable parameter, we computed Φ_MIP_ for *τ* from 1 (i.e., 1/20 s) to 100 (i.e., 5 s) frames. The time delay *τ* is the point at which the mean is the maximum 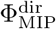 and 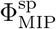 for all the datasets. Both peaks were located simultaneously. *τ* comprised approximately three frames (i.e., 0.15 s). This value was suitable because it was the same as the *ayu*’s response time (Figure S5).

#### Basic interpretation of IIT 2.0 in the group dynamics

Let us refocus on fish schools from perspective of the IIT. In group behaviour, the IIT describes an agent’s connectedness as a group. If all fish move randomly, Φ_MIP_ is expected to be zero because removing one fish does not affect the group’s behaviour. Similarly, if all fish move in the same direction and at the same speed, Φ_MIP_ becomes zero. Φ_MIP_ = 0 indicates that the group can be divided into two independent subgroups by MIP (Figure 1B). By contrast, the system must incorporate heterogeneous interactions to achieve a high Φ_MIP_. From an intrinsic perspective, the ‘differences that make a difference’ information matters. Therefore, highly integrated collective behaviour is neither ordered nor disordered. This is referred to as Φ_MIP_ of the ‘group integrity’. Φ_MIP_ evaluates the complex but incomparable group dynamics.

The main complex in the group corresponds to the core of information processing in the group (Figure 1B). In general, the main complex *M* is an appropriate subset of the entire set *S*. Because the highest Φ is related to the critical state (Figure S1; we show that a normal critical state, such as the SPP, has no parts for group criticality), this small group is expected to be highly susceptible to external perturbations. The interactions between these mutually susceptible subgroups determine how the main complexes are allocated to the entire group. Such a separation is based on the unique methodology of the IIT, wherein the theory does not discriminate between internal and external fluctuations.

Furthermore, we applied two Φs considering different information perspectives: orientation (i.e., 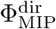 ) and speed (i.e., 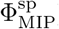 ), for two reasons: First, the criticality of orientation and speed is observed in animal groups [12]; however, the relation between these two critical dynamics is still uncertain. Examining group integrity from both perspectives facilitated the exploration of the relationship between these two criticalities from an IIT perspective. Second, the group requires a speed parameter to achieve group formation (e.g. schooling, milling, and swarming). We examined how these two Φ related to group formation (Figures S3 and S4).

#### Two types of integrity 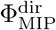 and 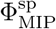

Figure 2A shows the time series of 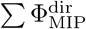 and 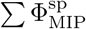 . Both values exhibit dynamic changes over time. Figure 2B shows the correlation between 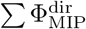 and 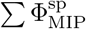 for all data. Although we observe a weak correlation between the two integrities, a consistent information flow between them cannot be confirmed (Table S1. However, upon increasing *T*_max_, we observe a statistically significant information flow). Therefore, the relationship between these two information processes can be considered independent.

**Figure 2:**
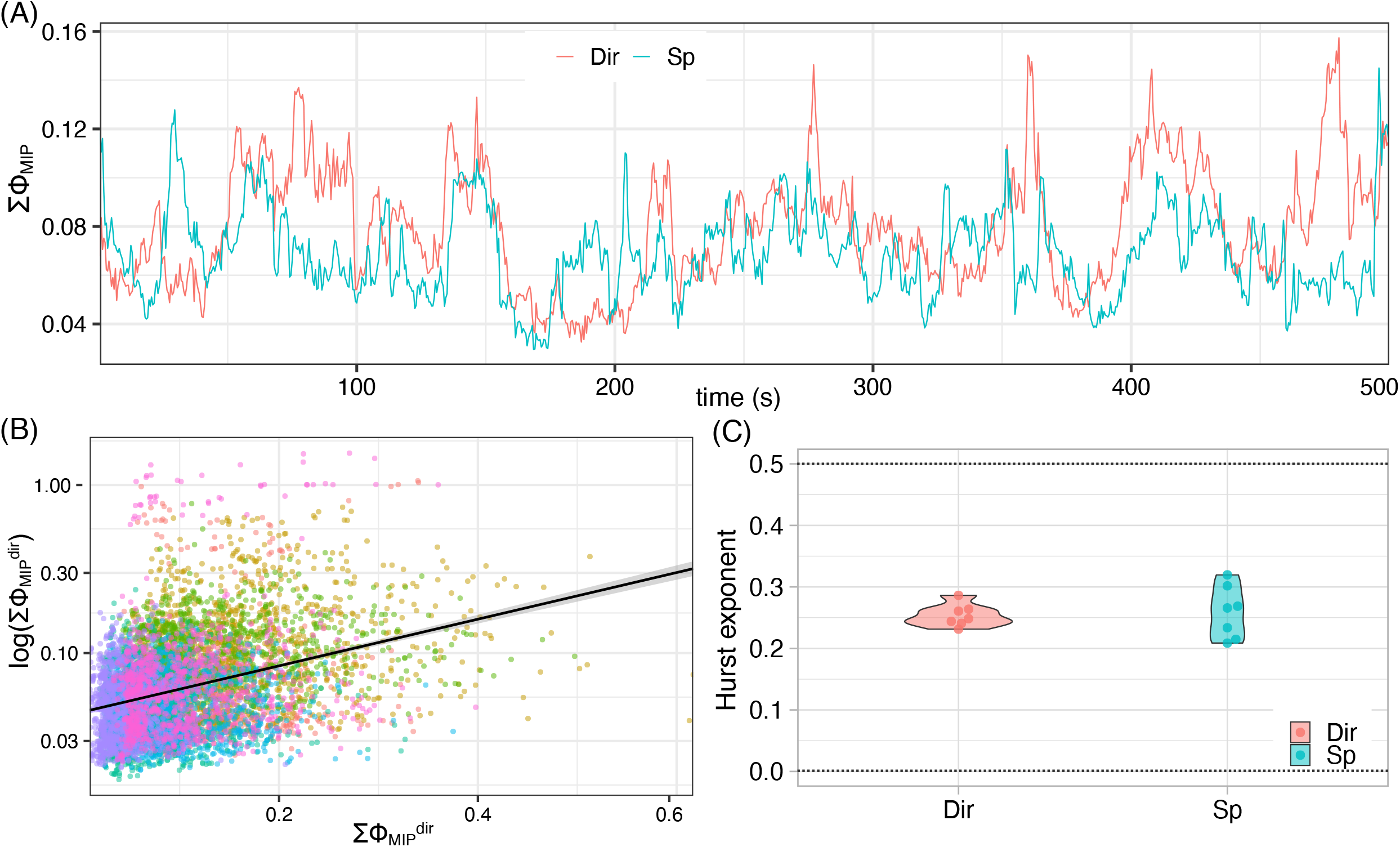
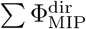 and 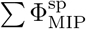 from real fish data. (A) Time series of 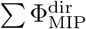 and 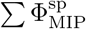. (B) Correlation between 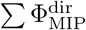 and 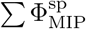 . Each colour corresponds to the data set number. (C) Violin plot of Hurst exponent (*H*) 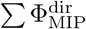 and 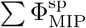. There is no significant difference between them.

However, the two information processes also shared a common ground. The most characteristic of these two time series was self-similarity (i.e. non-Brownian); the time series was affected by the past. Figure 2C shows the generalised Hurst exponent (*H*). If the time series is Brownian, the exponent *H* is approximately 0.5. If the time series is white noise, the exponent *H* is approximately zero. The exponent of the real-time series of both datasets is approximately 0.25 (Figure 2C). This self-similarity is also observed in the power spectrum analysis (Table S2). In contrast, the time series of max {Φ_MIP_ } is Brownian (Table 1). This difference implies that the real fish school has a historical effect as a whole (i.e. a set of all main complexes) but not for a certain subgroup as the maximum main complex.

**Table 1:**
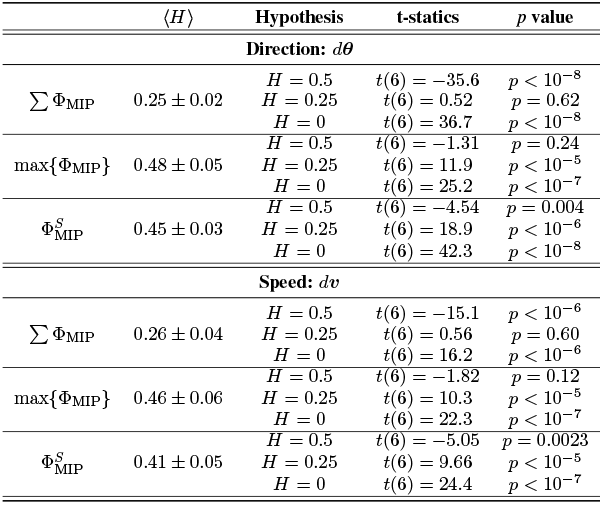
Statistical tests for ⟨*H*⟩ for each hypothesis. Note that *H* = 0 means the white noise and *H* = 0.5 indicates the Brownian noise (see Table S2 for the power spectrum analysis.)

#### Difference between the models and the real fish in terms of the integrity

Figure 3A shows the frequency distribution of the maximum main complex (MMC) size (i.e., |*M* |, where 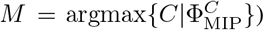 in the direction, speed, and SPP (10 agents). As listed in Figure S1, 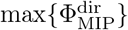 of the SPP model was located near the critical temperature. This result suggested the validity of our selection of the turning rate ***dθ*** as an IIT variable and that max{Φ_MIP_} could be a measure of the degree of criticality in SPP. Furthermore, a stark contrast was observed between the actual fish data and SPP model. The MMC size of the SPP was not fragmented (10/10 in most cases). Considering the distribution of the SPP exacted from the critical state, the critical state of the SPP indicated the indecomposable (non-divided) system from the IIT perspective. This result matched the critical-state property.

**Figure 3:**
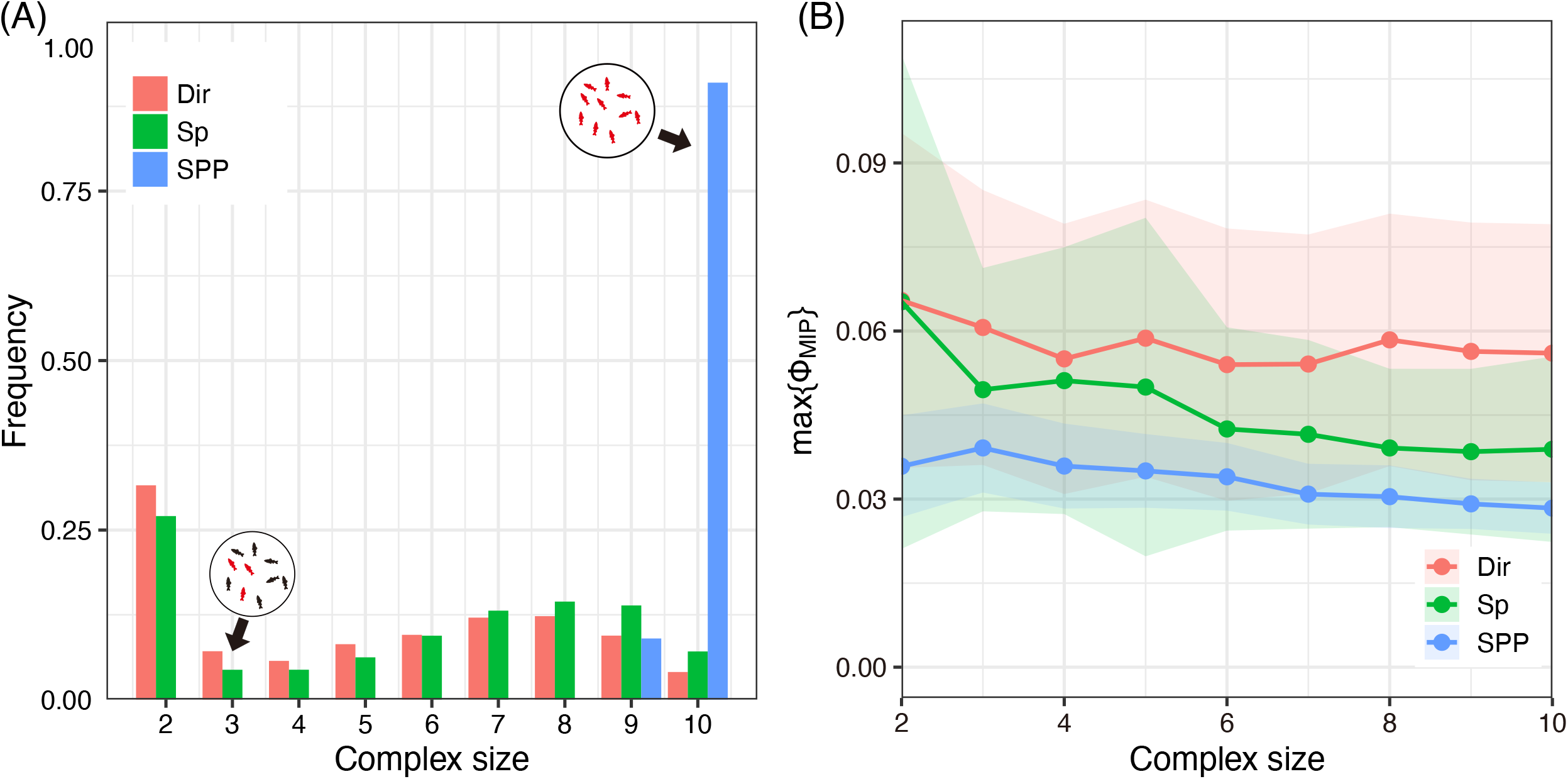
Data comparison with SPP. (A) Frequency distribution of the MMC for the direction (red), speed (green), and SPP(blue). The group size is 10. Inbox figures represent an example of MMC distribution (coloured red). (B) Mean max{Φ_MIP_} for each MMC size (the direction: red, speed: green, and SPP: blue).

In contrast, the MMC size of the real fish data was widely fragmented. The MMC sizes ranged from 2 (minimum) to 10 (maximum). Figure 3B shows the max{Φ_MIP_} for each MMC. All 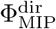 values of each MMC were significantly larger than those of SPPs. This result suggests that each fragmented MMC of real fish exhibited a critical state. Although our analysis did not prove that the high 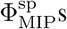s were also critical states (in the original SPP model, the velocity was set to be constant), our results suggest the coexistence of several degrees of critical states (i.e. measured by Φ_MIP_) at various levels (i.e. subgroups and behaviours). The critical state observed in real fish may not be a theoretically homogeneous phenomenon, similar to an SPP but heterogeneous.

Interestingly, we also found that 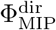of an entire set *S* for all samples (note that set *S* is not necessarily the main complex) still exhibited larger values than those of the SPP critical states 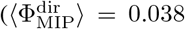. Welch t-test: *t*(148.6) = −19.2, *p <* 10^−30^). The complete integrity 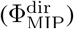 also exhibited criticality (the time series of 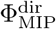 for the entire set *S* was not Brownian, as in Table 1). The high integrity Φ^dir^ of the entire group size was consistent with the classical result, which states that the entire group fluctuation was in a critical state. Thus, heterogeneous and homogeneous criticalities can coexist.

Next, we examine the relationship between max {Φ_MIP_}and the group formation. Generally, the Boid model can produce various group formations (e.g. swarms, milling, and schooling). The following parameters define the group formation:

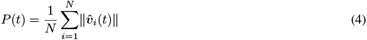

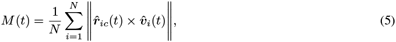

where 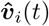 is the unit velocity vector and 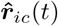 is the unit relative position vector from the centre of mass. *P* (i.e., polarity) measures the degree of group alignment (i.e., school formation). *M* (i.e., torus) measures the degree of group milling. *P* and *M* do not simultaneously have high values. Swarming formation corresponds to low *P* and *M* .

We found that the peaks 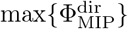 and 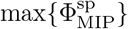 never overlapped in the Boid model (Figures S3 and S4). The peak of 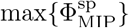 occurred during milling, and that of 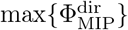 occurred within the transition area between schooling and swarming. From the perspective of the Boid-type model, group formations (*P* and *M* ) and group integrities (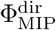 and 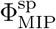 ) appeared to exhibit an intimate relationship.

However, Figure 4 in a real fish school revealed a different story. The max{Φ_MIP_} values for both cases correlated (see the Supporting Information for different values: Figure S6 and Figure S7). Furthermore, the 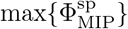 distribution was significantly different from the Boid distribution. In contrast to the Boid distribution, the speed integrity exhibited a high 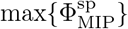 around the high-alignment area (i.e., ⟨*P* ⟩ ≈ 1). These results suggest that the information structure of a real fish school cannot be reproduced using a simple Boid-type model.

**Figure 4:**
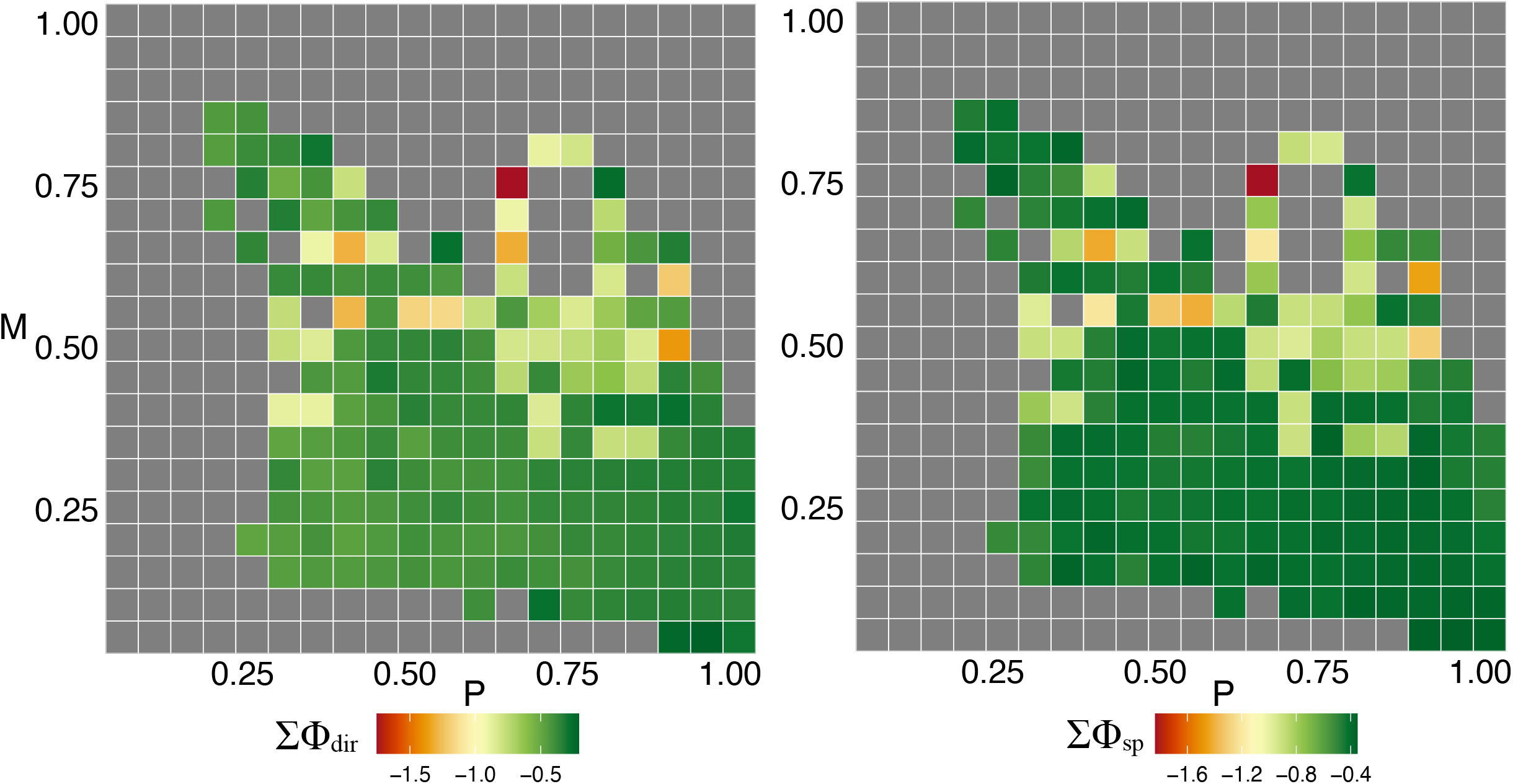
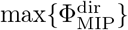 and 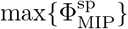 with the group formations (*P* and *M* ). The colour ba r represents the log-scale max {Φ_MIP_ }. *P* is a polarity and *M* is a milling for each data. We listed other heatmaps ΣΦ_MIP_ (Figure for S6) and complex size (Figure S7).

#### Role of MMC individuals

Because we found that the time series of 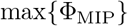 was Brownian, we examined whether the survival time of MMC was also a random process. The survival time for each individual was defined. The survival distribution can be approximated using a cumulative Weibull distribution, as follows:

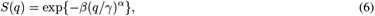

where *γ* is the average lifespan and *β* and *α* are the parameters to be estimated. *α* determines the shape of the exponential distribution. If *α*≈ 1, then the decay is exponential (i.e., there is no historical effect on MMC survival). In contrast, if *α <* 1, the graph is called ‘stretched exponential’ (i.e., the long-tail distribution but not the power law distribution).

Figure 5 shows the lifespan distribution for the real and shuffled fish data (an average of 100 samples). The shuffled distribution had no memory effect (i.e., 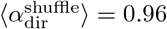 and 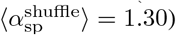). By contrast, the real fish distributions were highly stretched in both cases ( ⟨*α*_dir_⟩ = 0.61 ± 0.03 and ⟨*α*_sp_⟩ = 0.61 ± 0.02). Therefore, although max Φ_MIP_ was a Brownian process, the MMC series was not random, and it exhibited a highly heterogeneous distribution (see other results in Table S3).

**Figure 5:**
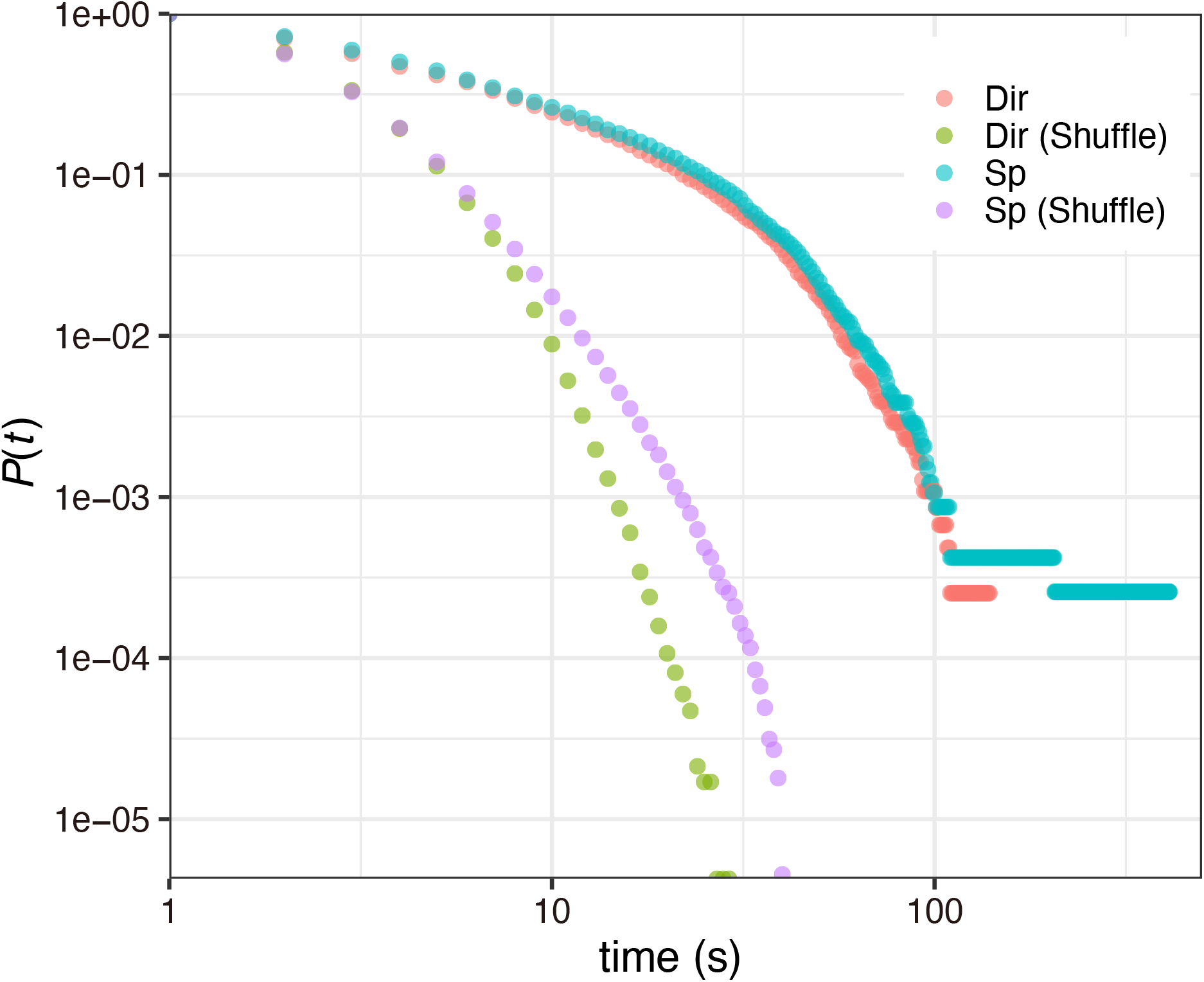
Survival probability *P* (*t*) of the MMC lifespan.

The heterogeneity of MMC lifespan suggests that being an MMC member for each fish may not be uniform. Certain members were highly allocated to the MMC, whereas others were not. The MMC rate is defined as follows:

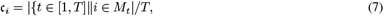

where *M*_*t*_ is the MMC at time *t* and *T* is the entire time of the data series. We refer to c_*i*_ as the ‘core rate’ for fish *i*. Figure 6A shows the core rate for direction 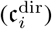 and the logarithm of the standard deviation of *dθ*_*i*_ (*dv*_*i*_) through the entire data series (no correlation was observed with mean *dθ*_*i*_: *n* = 70, *r* = 0.104, *p* = 0.392. In contrast, the mean *dv*_*i*_ was highly correlated with 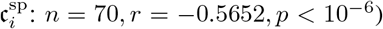. This negative correlation suggests that the fish belonging to the MMC group moved more stably (i.e., less affected by others’ movement) compared with the low-frequency fish. Thus, most individual activities in fish schools were too noisy for criticality. To achieve a high group performance, the group requires unaffected individuals (i.e., low *SD*(*dθ*_*i*_) and *SD*(*dv*_*i*_)) to connect various demands from other fish. This characteristic is different from leadership, as its position never places the group head in the moving direction (see Movie S1).

**Figure 6:**
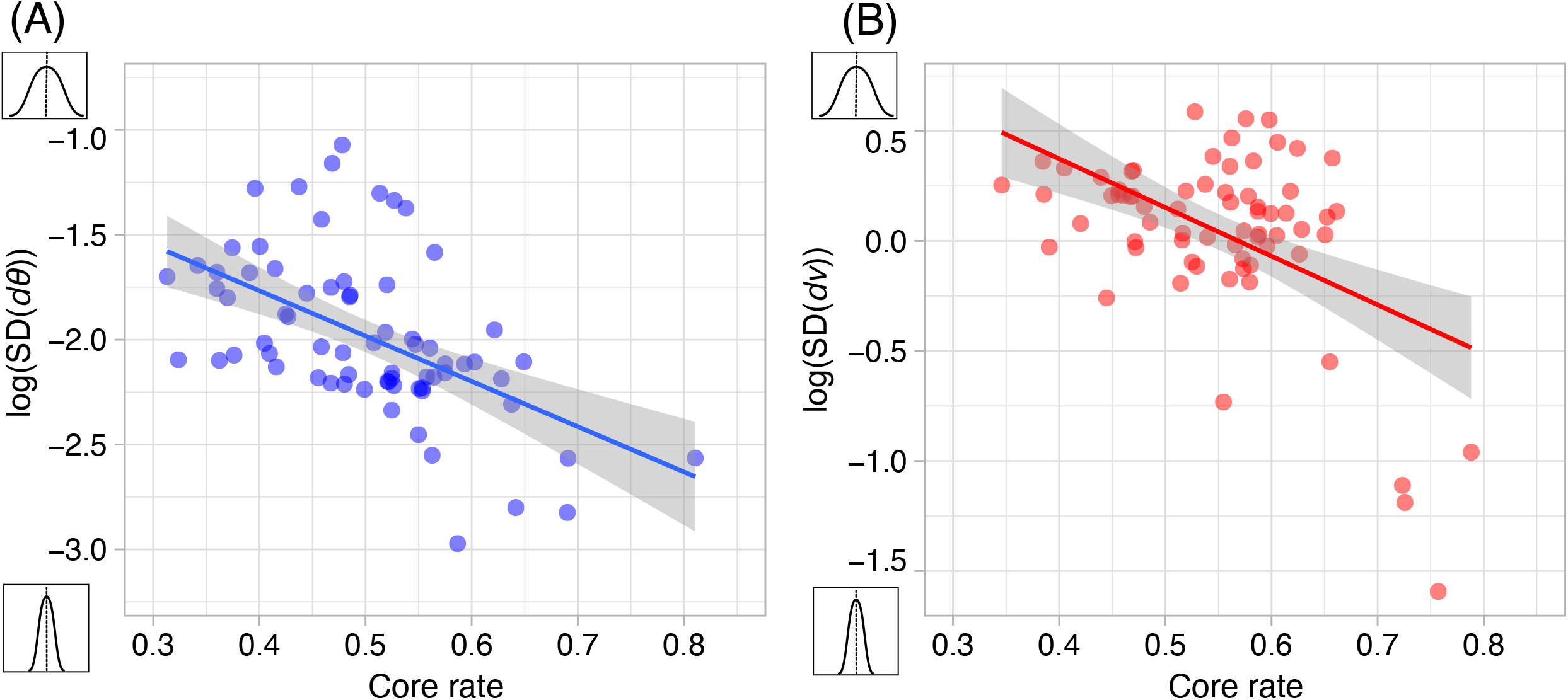
Correlation between the core rate (c_*i*_) and fish movements. (A) The direction deviation and core rate for direction (Pearson’s correlation test: *n* = 70, *r* = −0.536, *p <* 10^−6^). (B) Mean speed and the core rate for speed (Pearson’s correlation test: *n* = 70, *r* = −0.516, *p <* 10^−6^). Table S4 shows the results for other parameter settings.

## Discussion

In this study, we applied the IIT to an actual fish school (i.e. *ayu*s) and two representative models (SPP and Boid). We selected two variables (i.e., ***dθ*** and ***dv***) for the IIT analysis. Throughout the SPP analysis, the peak of the group integrity Φ_MIP_ corresponded to the phase transition point. This result also matched the previous IIT analysis of other criticality systems, which stated that Φ_MIP_ can measure the degree of criticality of the system.

Having identified the meaning of group integrity compared with the SPP, we found that the MMC of a real fish school comprised proper subsets. This result indicates that many degrees of criticality (e.g., intensity Φ_MIP_ and the size of group integrity |*M* | ) existed inside the group. This criticism provides a new perspective on traditional criticality arguments that state that a critical system is an inseparable system. Note that our argument does not contradict classical criticality. Instead, our findings indicate that global critical states (i.e., the entire group *S*) and local critical states (i.e., the subgroup *M* ) can coexist.

We found no evidence that the two groups’ integrities (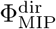 and 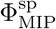 ) were related. Although we also observed the same type of self-similarity and a positive correlation between the two Φ_MIP_, there was no significant information transfer (i.e., transfer entropy) between them. This independence may originate from our definition of the group integrity of speed. Peak 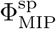 corresponded to a particular state (i.e., milling), whereas peak 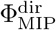 corresponded to the transition state. However, there may still be more appropriate variables for speed integration. We leave this issue for future research.

Furthermore, by examining the temporally heterogeneous MMC distribution, we found that high-frequency MMC members had small variances in direction and velocity. Our analysis suggested that lower-MMC members behaved more randomly than higher-MMC members (in terms of direction and speed). Regarding critical phenomena, our results suggest that global criticality can be divided into two subgroups: more affected individuals (i.e., quickly responding to internal/external perturbations) and less affected individuals (i.e., weakly responding to internal/external perturbations).

Global criticality may be a mixture of the different roles of groups. One interpretation of these role divisions within a group is as an adiabatic process [54]. Less affected individuals showed a slower response than more affected individuals. As Harken suggests, slow-response individuals enslave quick-response individuals [55]. In other words, multi-timescale interactions occur in collective behaviour [56]. Certain theoretical models support this consideration (e.g. the mutual anticipation of different timescales can enhance group dynamics [57, 58, 59]). The coexistence of several degrees of criticality may provide a key to understanding multi-timescale interactions.

However, certain problems must still be solved, such as the validity of the correspondence between a high Φ_MIP_ and critical state. In this study, we applied this correspondence to SPP results. However, the information process of the SPP is homogeneous (Φ_MIP_ does not depend on *T*_max_). Real fish schools are heterogeneous and (Φ_MIP_ varies depending on *T*_max_). The relatively heterogeneous process of the Boid model also exhibits a high Φ_MIP_ in the transition phase; however, the high Φ_MIP_ originates from the frequent fission–fusion process in the periodic boundary. The fish schools in our data were rarely split. Therefore, we cannot directly identify the high Φ_MIP_ of real fish using the modelling results. This problem does not arise from our method but from the fundamental difference between theoretical and empirical criticality, because nature has no theoretical criticality. Our study offers a possible approach for uncovering the detailed dynamics of critical phenomena in living systems. Based on our findings, we should examine these relationship in more detail to bridge the gap between theoretical and empirical critical phenomena.

## Methods

### Ethics statement

This study was conducted in strict accordance with the recommendations of the Guide for the Care and Use of Laboratory Animals from the National Institute of Health. The study protocol was approved by the Committee on the Ethics of Animal Experiments at the University of Tsukuba (Permit Number: 14-386). All efforts were made to minimise animal suffering.

### Experimental settings

We studied *ayu*s (*Plecoglossus altivelis*), also known as sweetfish, which are found throughout Japan and are widely farmed. Juvenile ayus (approximately 7–14 cm in body length) display typical schooling behaviour, although adult ayus tend to exhibit territorial behaviour in environments with low fish density. Juveniles purchased from Tarumiyoushoku (Kasumigaura, Ibaraki, Japan) were housed in a controlled laboratory. Approximately 150 fish lived in a 0.8 m3 tank of continuously filtered and recycled fresh water with a temperature maintained at 16.4 °C, and they were fed commercial food pellets. Immediately before each experiment, randomly chosen fish were separated to form a school of each size and moved to the experimental arena without pre-training. The experimental arena comprised a 33*m*^2^ shallow white tank. The water depth was approximately 15 cm; therefore, the schools were approximately two-dimensional. The fish were recorded with an overhead grayscale video camera (Library GE 60; Library Co. Ltd., Tokyo, Japan) at a spatial resolution of 640 × 480 pixels and a temporal resolution of 100 frames per second.

### Tracking System

We tracked the trajectories of *ayu* fish schools with *N* = 10 using seven samples (8–12 min recording length). We used the semi-auto tracking system (Move2D; Library GE 60; Library Co. Ltd., Tokyo, Japan) to obtain positional information. All the occlusion events were processed manually. Each frame was set to 1/20 s.

### Data treatment for the IIT application

The fish trajectory data were smoothed before obtaining the orientation and speed from the raw positional data. The numpy.convolve function was applied to the trajectory data for one frame (1/20 s) to remove noise.

## Acknowledgments

Support for data collection was provided by a Grant-in-Aid for Scientific Research from the Japan Society for the Promotion of Science (21H05302 to T.N.). The funders had no role in the study design, data collection and analysis, decision to publish, or manuscript preparation.

## Data availability

All data are available from https://github.com/t-niizato/Information-structure-of-heterogeneous-criticality.

### Competing interests

The authors declare no competing interests.

## Notes

https://github.com/t-niizato/Information-structure-of-heterogeneous-criticality

### Competing Interest Statement

The authors have declared no competing interest.

