## Supporting Information for "Information structure of heterogeneous criticality in a fish school"

#### Self-Propelled Particle (SPP) model on IIT

The self-propelled particle (SPP) model is widely used to study collective behaviour [1]. In particular, phase transitions in the SPP model have been discussed in various contexts. Recently, researchers have suggested that integrated information theory (IIT) can be applied to measure the degree of criticality [2, 3]. For instance, Mori et al. suggested that $\Phi_{MIP}$ in IIT 2.0 can estimate the phase transition area more accurately than mutual information [2]. However, there have been no reports on whether IIT can be used to estimate the phase transition points of an SPP.

First, we briefly provide an overview of the SPP. The agents of SPP are randomly distributed and have random directions (we considered only two dimensions). Each agent can change its direction according to its neighbour's direction within a fixed radius. There is one noise factor in the model. The noise term was added to the determined direction after determining the agent's next direction (the average of its neighbour's direction, including self). The equation used is as follows:

$$\theta_i(t+1) = \langle \theta(t) \rangle_r + \Delta\theta \quad (1)$$

where $\theta_i(t)$ is the direction at time $t$, $\langle \theta(t) \rangle_r$ is the mean direction of its neighbours within radius $r$, and $\Delta\theta$ is the noise term, which ranges from $[-\eta/2, \eta/2]$. $\eta$ is the noise parameter of the model and varies between 0 and $2\pi$. After determining the directions of all the agents, each position $\boldsymbol{x}_i(t)$ is updated simultaneously.

$$\boldsymbol{x}_i(t+1) = \boldsymbol{x}_i(t) + \boldsymbol{v}_i(t)\Delta t \quad (2)$$

where $\boldsymbol{v}_i$ is the direction vector and $\Delta t$ is the time step (defined $\Delta t = 1$). In general, this boundary condition is preferred because group cohesion can be maintained.

Vicseck et al. showed that a phase transition occurs from an ordered state to a disordered state. The agent direction was highly aligned for the lower temperature regions (i.e. low-noise parameters). In contrast, there was no such alignment in high-temperature regions (i.e. high-noise parameters), and all directions are randomly distributed. The critical state refers to the point at which the rotational symmetry breaks. The order parameter is expressed mathematically as follows:

$$P(t) = \frac{1}{N}\sum_{i=1}^{N}\left\|\frac{\boldsymbol{v}_i(t)}{\|\boldsymbol{v}_i(t)\|}\right\| \tag{3}$$

where $\|\boldsymbol{v}\|$ indicates the norm of vector $\boldsymbol{v}$. The order parameter $P$ varies from 0 (i.e. a disordered state) to 1 (i.e. an ordered state). The susceptibility $\xi = \langle P^2 - \langle P^2 \rangle\rangle$ reaches its maximum at the critical noise level. This value changes with the system size. In general, a well-established criticality estimation method is the finite-size scaling method.

In this section, we estimate the SPP criticality via the IIT. To achieve this, we must select the appropriate variables. In this study, we selected $d\theta$ as the turning rate from the previous direction. We chose this value because the direction change plays an essential role in the order parameter. Therefore, the data at time $t$ represent $[d\theta_1, d\theta_2, ...d\theta_N]$. As noted in the main text, we must also select time delay $\tau$. We set $\tau$=1 owing to the absence of information delay in this system.

Figure S1A shows the computational results from $N$=2 to $N$=10 in the SPP model. All cases indicate that the lowest $\Phi_{\mathrm{MIP}}$ is in the high-noise parameter region. This tendency makes sense of the IIT concept. Because the directions of all agents tend to behave randomly in this region, we cannot expect any integrity in this state. However, relatively high values were observed in the low-noise parameter region. Furthermore, the peak $\Phi_{\mathrm{MIP}}$ is located around the phase transition point. We computed the critical noise using finite-size scaling (we estimated $\eta \approx 2.57$ rad).

Figure S1B presents the average size of the main complex for each noise parameter. The size of the main complex exhibited a maximum rate below the critical point. From the IIT perspective, this result indicates that the system cannot be divided into two parts before the critical noise. Simultaneously, the dividing cost of the system reached a maximum near the critical point (Fig S1B). This observation aligns with the high susceptibility of the system at the critical point because the long-range correlation indicates the degree of connectedness as one system (i.e., the system is difficult to separate). Therefore, the IIT can be a good measure of the degree of criticality of collective behaviour.

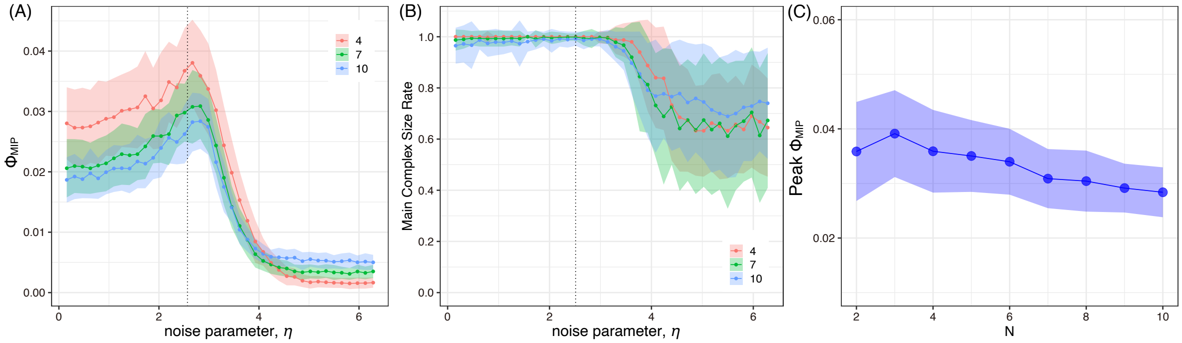


Figure S1: $\Phi_{MIP}$ **with noise parameter (left), the main complex rate with noise parameter (middle), the maximum** $\Phi_{MIP}$ **value for each group size**(A) $\Phi_{MIP}$ peaks at the 2.57 (rad) for each group size. (B) Main complex rates are the maximum in the low-noise region. (C) Maximum $\Phi_{MIP}$ for each group size is selected. $\Phi_{MIP}$ gradually decreases with the group size. We set the $T_{max}$ = 500 as the same as the real fish analysis. If we change $T_{max}$ = 10000, the peak never changes.

However, the highest $\Phi_{MIP}$ value reflected a critical point. We assumed a high density ($L^2/N = 4$). The agent's interaction range overlapped with that of the other agents. We consider this assumption valid based on our experimental data; however, to evaluate the precise meaning of IIT, the low-density case must also be examined.

Figure S2A shows $\Phi_{MIP}$ for the low-density conditions. $\Phi_{MIP}$ values exhibited relatively low values at low noise levels. This result arose from the fact that the agent sometimes moved separately because of the vast space in its interaction domain. Interestingly, in contrast to the dense SPP results, the $\Phi_{MIP}$ peak shifted to a lower noise parameter as the group size increased. This peak shift can be inferred from Figure S2B. In low-noise-parameter regions, the maximum complex becomes fragmented. This decreased MMC size suggested that the groups were not connected to each other because of the low-density condition. As observed in the SPP, the interaction among the agents was considered the collision of the fragmented group. Therefore, large fluctuations occurred in the low-noise parameter regions. Note that this high group integrity $\Phi_{MIP}$ was accomplished by a periodic condition. Therefore, we must distinguish this low-density behaviour from the high-density behaviour.

Notably, the behaviour of the high-density groups exhibited a consistent trend concerning IIT, specifically regarding the peak position of $\Phi_{MIP}$ and the corruption position of the complex size rate, regardless of the group size. The IIT can estimate the critical point in this homogeneous system without using the finite-size scaling method; however, this issue is beyond the scope of this study and has been left to future research. Herein, we confirm that $\Phi_{MIP}$ and the complex size can measure the degree of criticality in collective behaviour.

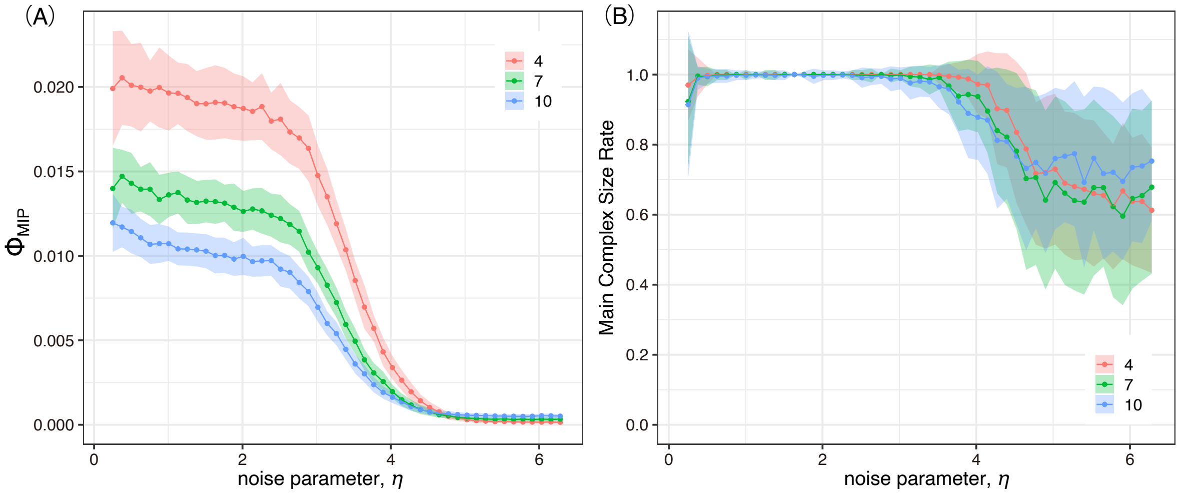


Figure S2: $\Phi_{\mathrm{MIP}}$ **with noise parameter (left), the main complex rate with noise parameter (right).** (A) $\Phi_{\mathrm{MIP}}$ and the noise parameter at low density. (B) Main complex rates are the maximum in the low-noise region. We set the $T_{\max} = 10000$ steps to obtain stable results.

### BOID model on IIT

Boid is the most popular model of collective behaviour. A Boid-type model was constructed using repulsion, alignment, and attraction. Each agent had three distinct interaction regions allocated to three separate interactions.

The interaction in the alignment region was identical to that in the SPP interaction. In the attraction regions, the agent directed the centre of mass to the mass of its neighbours. In contrast, the agent indicated the opposite direction to the central neighbour position in the repulsion regions. In general, the Boid model combines these three interactions.

In this study, we applied Couzin's model [4] for our analysis because their model is the archetype of the present collective behaviour model. The model required various optional parameters, such as visual fields ($\alpha = 1.5\pi$ (rad)), turning rate ($d\theta_{\max} = 0.7$ (rad/s)), speed range ($\|\boldsymbol{v}\| = 1 \sim 5$ (m/s)), and error rate ($\sigma = 0.05$ (rad)). We set the same parameters as those in our settings. In their study, the group formed various shapes (e.g., milling, schooling, and swarm) according to the weight of the alignment and attraction parameters.

This section analyses the IIT context. The merits of Boid analysis are twofold. First, Couzin's Boid model considers the speed tuning of an agent. Their model defines the speed of the agent as the norm of the velocity vector. Because the SPP model does not affect speed differences, Couzin's model can estimate the effect of speed control in a group on group integrity (i.e. $\Phi_{\mathrm{MIP}}$ and the main complex). Second, in actual collective behaviour, group shape is also important. We estimated the meanings of the various forms from the IIT perspective. For example, the milling form, which represents the torus structure, can be achieved using an optimal turning rate and speed variation. Our interest lines in the relation of two ($\Phi_{MIP}^{dir}$: direction and $\Phi_{MIP}^{sp}$: speed) under this formation.

The definition of $\Phi_{MIP}^{sp}$ is the same as that in the main text ($T_{max} = 500$). We computed all $\Phi$ values for the parameter sets (attraction and alignment parameters: repulsion was set at 0. 100 trials for each parameter set). Figure S3A and S3B show the mean polarity $\langle P \rangle$ and torus $\langle M \rangle$ for each parameter set, respectively. Torus was defined as follows:

$$M(t) = \frac{1}{N}\sum_{i=1}^{N} \|\hat{\boldsymbol{r}}_{ic}(t) \times \hat{\boldsymbol{v}}_i(t)\| \tag{3}$$

where $\hat{\boldsymbol{x}}$ means the unite vector of $\boldsymbol{x}$, and $\boldsymbol{r}_{ic}(t) = \boldsymbol{x}_i(t) - \boldsymbol{x}_C(t)$ ($\boldsymbol{x}_C$ is the centre of mass). The low-alignment and high-attraction regions tended to undergo milling (Figure S3B). By contrast, the group formed a high school alignment area (Figure S3A). Figure S3C and S3D show the averages $\Phi_{MIP}^{sp}$ and $\Phi_{MIP}^{dir}$ for each set of parameters, respectively. The graph showed that the MIP peak $\Phi_{MIP}^{sp}$ was located around the torus region (i.e., the milling formation).

However, the high $\Phi_{MIP}^{dir}$ regions did not overlap with the high $\Phi_{MIP}^{sp}$ regions. Speed group integrity also reflected the morphological structure of the group (i.e. milling); however, the peaks of $\Phi_{MIP}^{dir}$ were located around the boundary regions where schooling, millings, and swarming occurred. The high group integrity in this direction was derived from fission–fusion dynamics rather than group dynamics as a single unit. Under the periodic boundary condition, each agent could interact with previously separated agents after certain time steps. This reorganisation of the group tended to increase integrity; however, this type of high integrity was never directly linked to consistent cohesion in real fish schools.

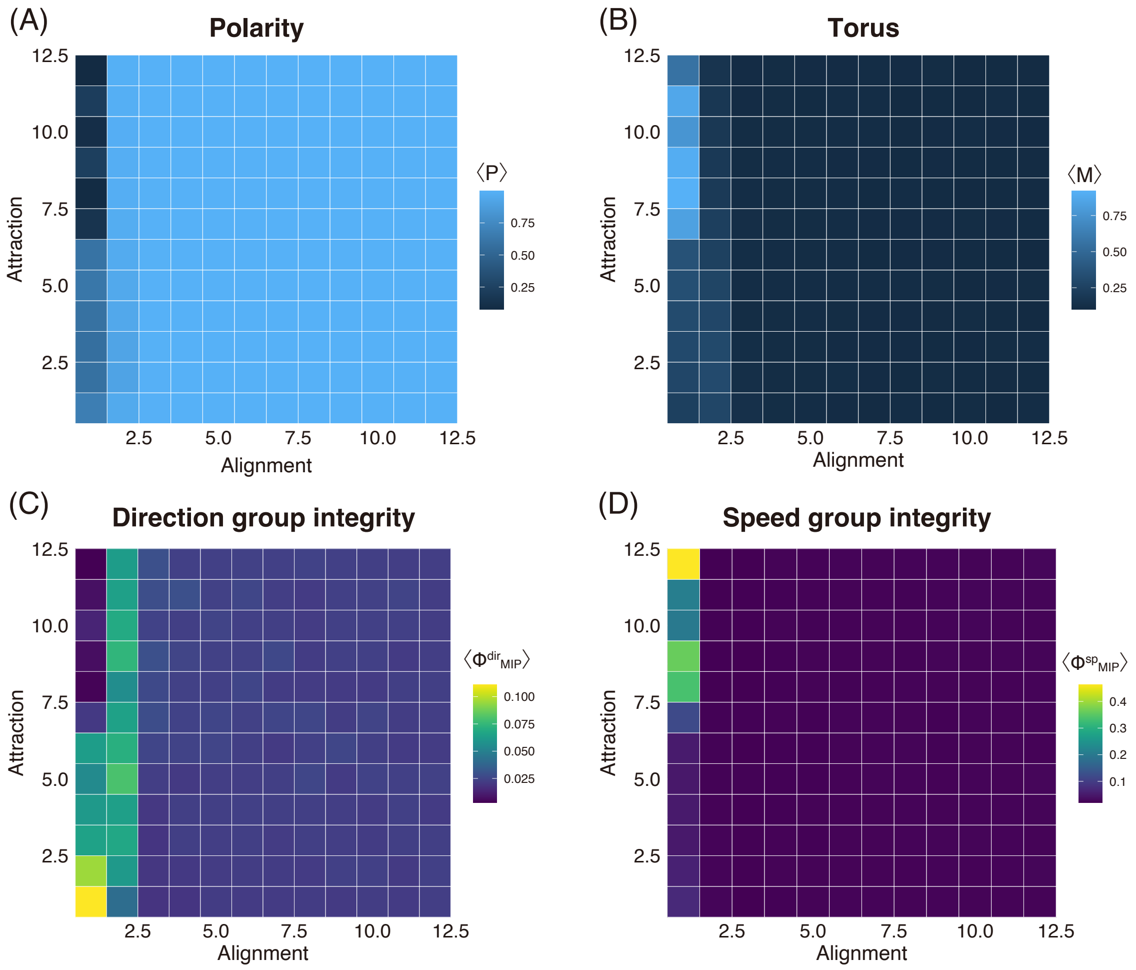


Figure S3: **Heatmap of ⟨P⟩, ⟨M⟩, $\Phi_{MIP}{}^{dir}$ and $\Phi_{MIP}{}^{sp}$.** (A, B) Group formation to the assigned parameter settings. (A) Polarity estimates the degree of alignment. (B) Torus estimates the degree of milling structure. (C, D) Group integrity to the assigned parameter settings. (C) $\Phi_{MIP}{}^{dir}$ for the same parameter setting. (D) $\Phi_{MIP}{}^{sp}$ for the same parameter setting.

Figure S4A shows the distribution of both averaged $\Phi_{MIP}$ values along the same horizontal axis, $\langle P \rangle$. $\Phi_{MIP}{}^{dir}$ (blue) and $\Phi_{MIP}{}^{sp}$ (red) are assigned different colours. The $\Phi_{MIP}{}^{dir}$ and $\Phi_{MIP}{}^{sp}$ peaks did not overlap, as shown in Figure S3 (CD). This tendency was the main characteristic of Boid-type interactions. The schooling formations exhibited no speed integrity. Torus formation showed no directional integrity. This result was natural, because one extreme obliged the other to hold a relatively uniform distribution to maintain a stable group shape (for instance, the high-alignment form requires a uniform speed distribution).

Figure S4B shows the main complex sizes of the direction and speed. As observed in Figure S4A, both main complex sizes do not overlap, except at extreme alignments (i.e., $\langle P \rangle \approx 1$).

However, the complete main complex around ⟨$P$⟩ ≈ 1 does not imply a high integrity. This tendency may be attributed to the uniform distribution of direction and speed (the integrity of both groups is approximately zero at ⟨$P$⟩ ≈ 1 in Figure S4A). The high $\Phi_{MIP}^{dir}$ values in the middle of ⟨$P$⟩ regions may be attributed to fission–fusion formation. Interestingly, the main complex size of the speed around the low- polarity region (i.e., the milling region) is fragmented. This fragmentation suggests the fragility of the milling structure in the general sense from an IIT perspective. In this sense, speed group integrity differs from direction group integrity.

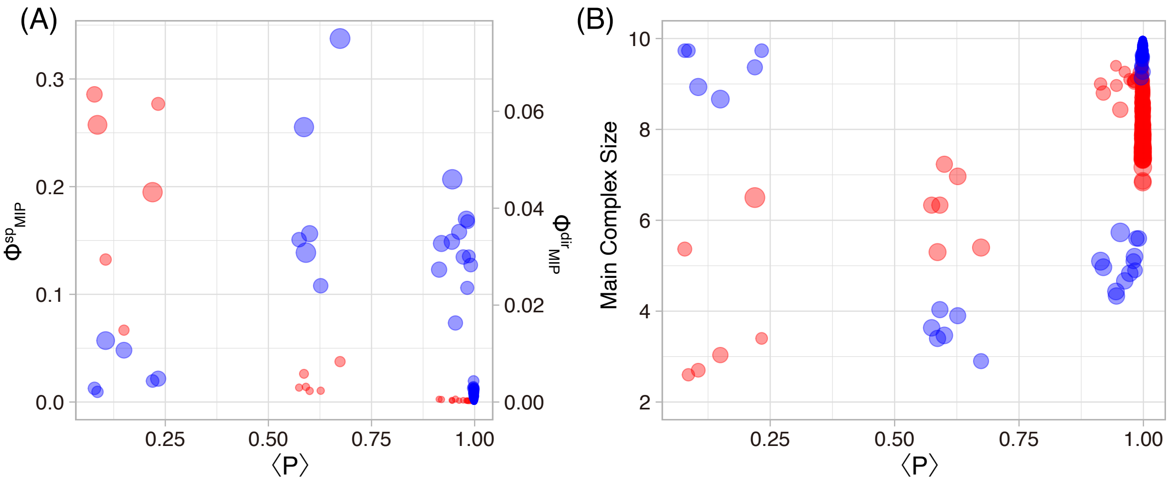


Figure S4. **IIT analysis for the Boid's flock**. (1) $\Phi_{MIP}^{dir}$ (blue) and $\Phi_{MIP}^{sp}$ (red) for the mean polarity ⟨$P$⟩. Each size of dot represents SD. (2) Main complex size of the direction and the speed for the mean polarity ⟨$P$⟩. Each size of dot represents SD.

[3] William Marshall, Hyunju Kim, Sara I Walker, Giulio Tononi, and Larissa Albantakis. How causal analysis can reveal autonomy in models of biological

systems. *Philosophical transactions of the Royal Society a: Mathematical, physical and engineering sciences*, 375(2109):20160358, 2017.

[4] Iain D Couzin, Jens Krause, Richard James, Graeme D Ruxton, and Nigel R Franks. Collective memory and spatial sorting in animal groups. *Journal of theoretical biology*, 218(1):1–11, 2002.

### Supporting Figures and Tables

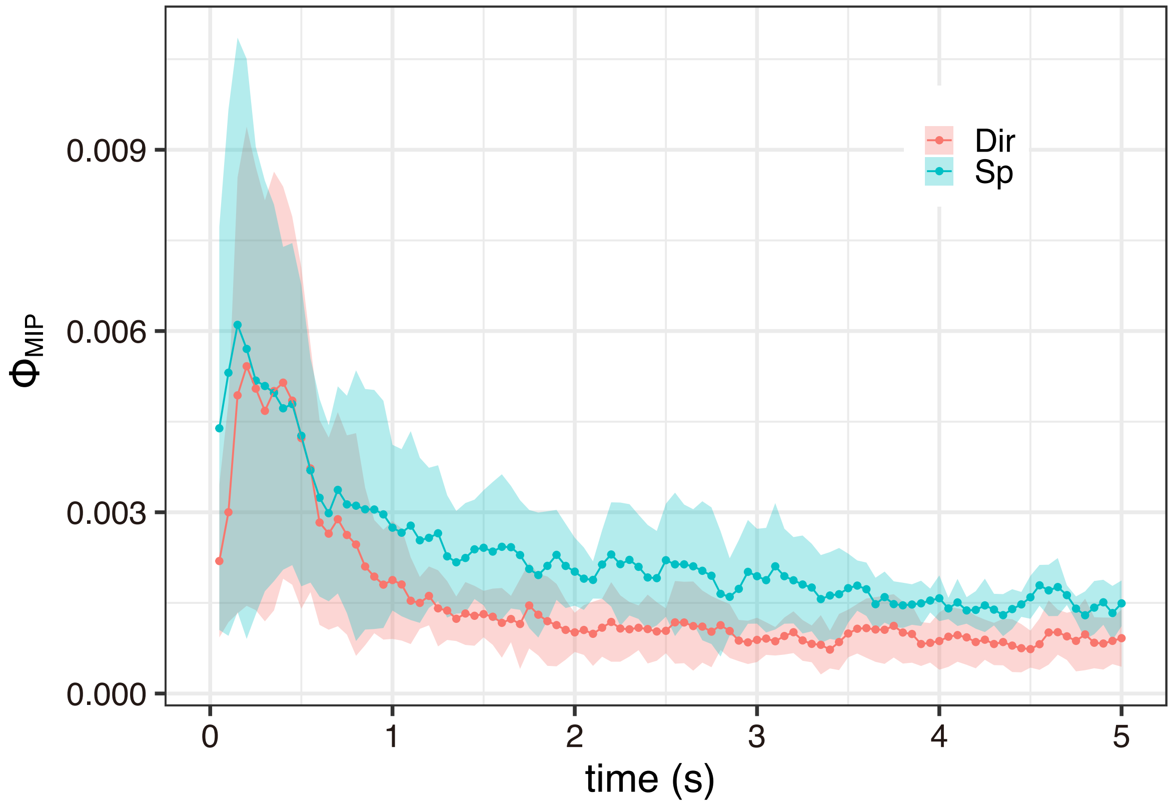


Figure S5: **Peak $\Phi_{MIP}$ for each $\tau$.** We select $\tau = 0.15$ as optimal.

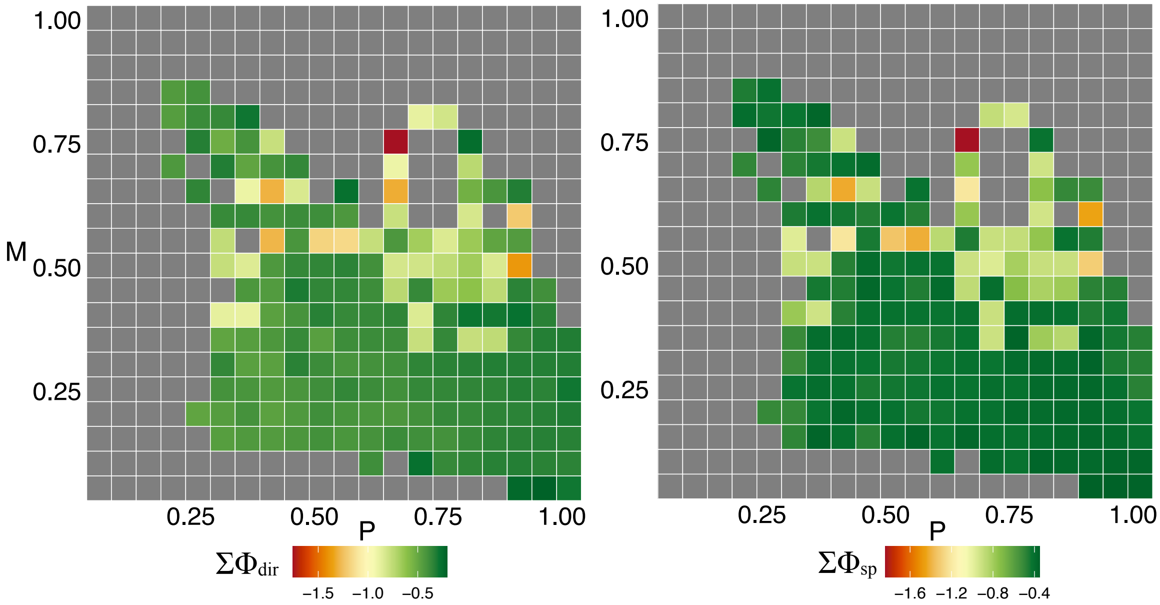


Figure S6: **$\Sigma\Phi_{MIP}{}^{dir}$ and $\Sigma\Phi_{MIP}{}^{sp}$ with the group formations** (*P* and *M*). The colour bar represents the log-scale $\Sigma\Phi_{MIP}$. *P* is a polarity and *M* is a milling for each data.

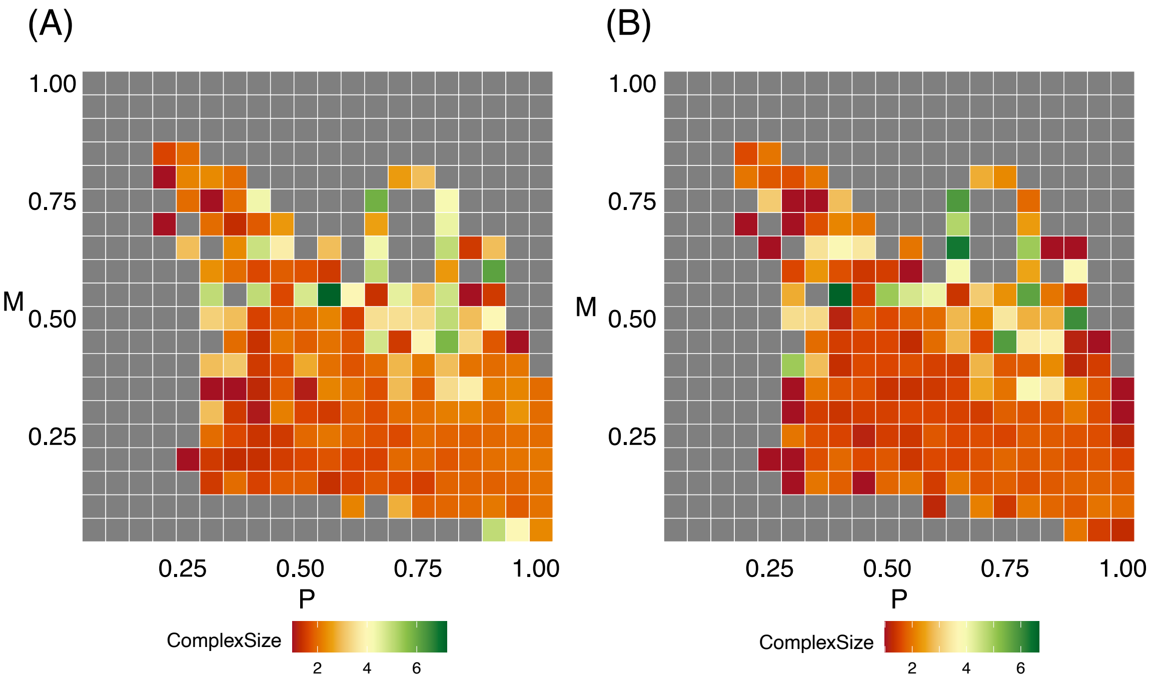


Figure S7: **Maximal main complex size for the direction and speed with the group formations** (*P* and *M*). The colour bar represents the MMC from 2 to 10. *P* is a polarity and *M* is a milling for each data.

Table S1: **Statistical test with transfer entropy between two group integrity for each $T_{max}$.**

| | L=200 | | | | |
|---|---|---|---|---|---|
| | | TR(Dir -> Sp) | p value | TR(Sp -> Dir) | p value |
| Σ Φ | 1 | 0.0065 | 0.46 | 0.0025 | 0.9633 |
| | 2 | 0.0051 | 0.54 | 0.003 | 0.8133 |
| | 3 | 0.0058 | 0.5433 | 0.0082 | 0.1767 |
| | 4 | 0.0056 | 0.2533 | 0.0043 | 0.61 |
| | 5 | 0.0053 | 0.4133 | 0.0016 | 0.9667 |
| | 6 | 0.0067 | 0.2667 | 0.0068 | 0.3267 |
| | 7 | 0.0111 | 0.0533 | 0.0171 | **0.0033** |
| | **average** | **0.0066** | | **0.0062** | |
| max{Φ} | 1 | 0.0066 | 0.2667 | 0.0118 | 0.0167 |
| | 2 | 0.0138 | **0** | 0.0039 | 0.71 |
| | 3 | 0.0087 | 0.1567 | 0.0093 | 0.07 |
| | 4 | 0.0048 | 0.44 | 0.0043 | 0.54 |
| | 5 | 0.0082 | 0.0833 | 0.0025 | 0.89 |
| | 6 | 0.0066 | 0.3367 | 0.0131 | **0** |
| | 7 | 0.0064 | 0.42 | 0.0129 | **0.0133** |
| | **average** | **0.0079** | | **0.0083** | |

| | L=400 | | | | |
|---|---|---|---|---|---|
| | | TR(Dir -> Sp) | p value | TR(Sp -> Dir) | p value |
| Σ Φ | 1 | 0.007 | 0.2233 | 0.0092 | 0.1233 |
| | 2 | 0.0039 | 0.7433 | 0.0094 | 0.0567 |
| | 3 | 0.0096 | 0.09 | 0.0059 | 0.5167 |
| | 4 | 0.0057 | 0.34 | 0.003 | 0.8467 |
| | 5 | 0.0041 | 0.67 | 0.0086 | 0.0567 |
| | 6 | 0.0055 | 0.46 | 0.0114 | **0.0067** |
| | 7 | 0.0053 | 0.35 | 0.0105 | **0.04** |
| | **average** | **0.0059** | | **0.0083** | |
| max{Φ} | 1 | 0.0022 | 0.8767 | 0.0092 | 0.16 |
| | 2 | 0.0056 | 0.2267 | 0.0058 | 0.2267 |
| | 3 | 0.0024 | 0.9767 | 0.011 | **0.03** |
| | 4 | 0.0062 | 0.1533 | 0.0039 | 0.6 |
| | 5 | 0.0052 | 0.3133 | 0.0033 | 0.6967 |
| | 6 | 0.0043 | 0.49 | 0.0024 | 0.86 |
| | 7 | 0.0033 | 0.54 | 0.0096 | 0.06 |
| | **average** | **0.0042** | | **0.0065** | |

| | L=600 | | | | |
|---|---|---|---|---|---|
| | | TR(Dir -> Sp) | p value | TR(Sp -> Dir) | p value |
| Σ Φ | 1 | 0.0057 | 0.4567 | 0.0079 | 0.1533 |
| | 2 | 0.0058 | 0.3833 | 0.005 | 0.5867 |
| | 3 | 0.0058 | 0.55 | 0.0087 | 0.1933 |
| | 4 | 0.0085 | **0.05** | 0.006 | 0.3067 |
| | 5 | 0.0121 | **0** | 0.013 | **0** |
| | 6 | 0.0112 | **0.01** | 0.009 | 0.0833 |
| | 7 | 0.0054 | 0.53 | 0.0061 | 0.4767 |
| | **average** | **0.0078** | | **0.0080** | |
| max{Φ} | 1 | 0.0166 | **0.01** | 0.0057 | 0.2767 |
| | 2 | 0.0036 | 0.6833 | 0.0049 | 0.41 |
| | 3 | 0.0118 | **0.02** | 0.0219 | **0** |
| | 4 | 0.0031 | 0.7867 | 0.0069 | 0.0967 |
| | 5 | 0.005 | 0.263 | 0.0051 | 0.253 |
| | 6 | 0.0117 | **0.003** | 0.0057 | 0.29 |
| | 7 | 0.0071 | 0.1833 | 0.0183 | **0** |
| | **average** | **0.0084** | | **0.0098** | |

Table S2: **The mean slop of $f^{-\alpha}$ between two group integrity for each $T_{max}$.** $\alpha \approx 1$ indicates the strong self-similarity.

| | | L=200 | | L=400 | | L=600 | |
|---|---|---|---|---|---|---|---|
| | | mean(α) | ST(α) | mean(α) | ST(α) | mean(α) | ST(α) |
| max{Φ} | direction | -1.488 | 0.027 | -1.651 | 0.139 | -1.666 | 0.099 |
| | speed | -1.467 | 0.128 | -1.652 | 0.103 | -1.688 | 0.175 |
| Σ Φ | direction | -0.841 | 0.083 | -0.965 | 0.045 | -0.979 | 0.044 |
| | speed | -0.856 | 0.108 | -0.997 | 0.180 | -1.070 | 0.166 |

Table S3: **The parameter of Weibull distribution for each $T_{max}$.**

| | L=200 | | L=400 | | L=600 | |
|---|---|---|---|---|---|---|
| | mean(α) | mean(β) | mean(α) | mean(β) | mean(α) | mean(β) |
| direction | 0.68 ± 0.02 | 1.25 ± 0.02 | 0.61 ± 0.03 | 1.31 ± 0.03 | 0.55 ± 0.04 | 1.36 ± 0.04 |
| speed | 0.68 ± 0.03 | 1.25 ± 0.04 | 0.61 ± 0.02 | 1.31 ± 0.02 | 0.58 ± 0.02 | 1.33 ± 0.03 |

Table S4: **The correlation between core rate ($c_i$) and fish movements for each $T_{max}$.**

| | L=200 | | | L=400 | | | L=600 | | |
|---|---|---|---|---|---|---|---|---|---|
| | data num | CC | p | data num | CC | p | data num | CC | p |
| Direction | 70 | -0.746 | **1.25E-13** | 70 | -0.536 | **1.72E-06** | 70 | -0.379 | **0.0012** |
| Speed | 70 | -0.702 | **1.30E-11** | 70 | -0.565 | **4.75E-06** | 70 | -0.412 | **3.93E-04** |
